# USPNet: unbiased organism-agnostic signal peptide predictor with deep protein language model

**DOI:** 10.1101/2021.11.04.467361

**Authors:** Shenyang Chen, Qingxiong Tan, Jingchen Li, Yu Li

## Abstract

Signal peptide is a short peptide located in the N-terminus of proteins. It plays an important role in targeting and transferring transmembrane proteins and secreted proteins to correct positions. Compared with traditional experimental methods to identify and discover signal peptides, the computational methods are faster and more efficient, which are more practical for the analysis of thousands or even millions of protein sequences in reality, especially for the metagenomic data. Therefore, computational tools are recently proposed to classify signal peptides and predict cleavage site positions, but most of them disregard the extreme data imbalance problem in these tasks. In addition, almost all these methods rely on additional group information of proteins to boost their performances, which, however, may not always be available. To deal with these issues, in this paper, we present Unbiased Organism-agnostic Signal Peptide Network (USPNet), a signal peptide prediction and cleavage site prediction model based on deep protein language model. We propose to use label distribution-aware margin (LDAM) loss and evolutionary scale modeling (ESM) embedding to handle data imbalance and object-dependence problems. Extensive experimental results demonstrate that the proposed method significantly outperforms all the previous methods on the classification performance. Additional study on the simulated metagenomic data further indicates that our model is a more universal and robust tool without dependency on additional group information of proteins, with the Matthews correlation coefficient improved by up to 17.5%. The proposed method will be potentially useful to discover new signal peptides from the abundant metagenomic data.

## 1 Introduction

A signal peptide (SP) is a short amino acid sequence working as a specific targeting signal to guide and transfer proteins into secretary pathways. It has a three-domain structure: Positively charged N-region, hydrophobic H-region and uncharged C-region [1]. The SPs function as specific segments to guide proteins to reach correct positions and then be cleaved by cleavage sites nearby its C region. Thus, the identification of signal peptides is vital for studying destinations and functions of proteins [2].

Many experimental and computational tools have been proposed to classify signal peptides and predict cleavage sites. The first attempt was a formulating rule proposed in 1983 [3]. Von Heijen firstly applies a statistical method to unveil patterns nearby cleavage sites of signal peptides based on only 78 Eukaryotic proteins [3]. Furthermore, generative model, such as hidden Markov model (HMM), are proposed to facilitate recognition of signal peptides. These models focus on analyzing these three functional regions (N-region, C-region, and H-region) in detail and are built by capturing the relationships between different regions of signal peptide [4, 5, 6, 7, 8, 9, 10]. Different from generative models, some homology-based methods are proposed [11]. The predictions of these methods are based on the similarities between sequences in existing knowledge base and the input sequences. In addition, they can achieve similar prediction performance as generative models.

Recently, supervised models make great progress in the recognition of signal peptides. The query sequences are encoded into embedding vectors and then fed into models to directly compute probabilities for each signal peptide type. Among these models, machine-learning based models take an important role for their remarkable performances. SignalP 4.0 proposed to use pure neural network architecture instead of generative models [12]. DeepSig applied deep convolutional neural networks (DCNNs) architecture to the recognition of signal peptides and the prediction of cleavage site positions [13]. Furthermore, SignalP 5.0 came up and benchmarked all the previous proposed methods [14]. These methods achieved advanced performance in tasks, but most of them suffer extreme class-imbalance and therefore perform poorly on minor classes. In addition, these methods often depend heavily on additional information about group of organisms to boost their performances. However, it is impractical to obtain sufficient group information from metagenomic data in reality. A robust tool should only require amino acid sequences to yield accurate prediction results.

In this paper, we focus on solving the problems related to imbalance of training data and object-dependence on group information in signal peptide prediction. The main contributions are summarized as follows:

- We are the first to resolve the existing extreme imbalance problem in signal peptide prediction. Considering that previous algorithms train models mainly based on cross entropy loss, we propose to apply label-distribution-aware-margin loss (LDAM) to improve generalization of less frequent classes [15]. Furthermore, we present a modified loss function by combining class-balance loss with LDAM loss to improve generalization It is inspired by the point proposed in LDAM Loss that the novel margin-based loss is orthogonal to reweighting techniques.
- We introduce evolutionary scale modeling (ESM) embedding to enrich our embedding [16]. The contextual language model is trained based on 86 billion amino acids by unsupervised learning. The structural representations extracted from the protein language model not only contribute to production of state-of-the-art features as part of embedding to further facilitate signal peptide prediction, but also can capture biological properties of amino acid sequences in deep and compensate for loss of group information, thus building up an organism-agnostic signal peptide predictor to classify unknown proteins groups.
- We propose a new deep learning model called Unbiased Organism-agnostic Signal Peptide Network (USP-Net). Extensive experiments demonstrate that the proposed method achieves the state-of-the-art performance over other signal peptide predictors on signal peptide classification, especially on the organism-agnostic signal peptide prediction. When tested on metagenomic data, the model reaches higher MCC compared with SignalP5.0, up to 17.5%.

## 2 Methods

### 2.1 Datasets

In this study, the benchmark datasets are composed of two components. The first one is collected from SignalP5.0, which relies on the UniProt Knowledgebase released in April 2018 [17]. The other dataset is an independent dataset, SP19, firstly proposed in SignalP3L 3.0, which collected proteins from Swiss-Prot Knowledgebase from April 2018 to July 2019. The original SignalP5.0 dataset is composed of proteins from four groups: Eukaryotes, Gram-positive, Gram-negative bacteria, and Archaea. With aim to verify the effects of better generalization on minor classes, we carry out three separate SP types: Sec/SPI, Sec/SPII, and Tat/SPI SPs. Other proteins are accordingly considered as TM/Globular (NO-SP) type. The dataset with 4 separate labels has a long-tailed label distribution, which means that NO-SP type is superior in numbers among different labels. We continue to follow the two-run mode test of SignalP5.0, which means that we will firstly consider only the relative SP type as “positive dataset” and all the TM/Globular protein as “negative dataset”. Then, the other two SP types are added into “negative dataset” for further run. Here, we take SignalP5.0 dataset as the basis for our training dataset and benchmark dataset. The details of the training dataset are provided in Table 1.

**Table 1:** Statistics of training dataset adopted in this study. T denotes Tat/SPI, L denotes Sec/SPII and N/C denotes TM/Globular(NO-SP) here.

| Dataset | Organism | SP | T | L | N/C | Total |
| --- | --- | --- | --- | --- | --- | --- |
| SignalP5.0 | Eukaryotes | 2614 | 0 | 0 | 14656 | 17270 |
|  | Gram-positive | 189 | 95 | 449 | 190 | 923 |
|  | Gram-negative | 509 | 334 | 1063 | 422 | 2328 |
|  | Archaea | 60 | 27 | 28 | 122 | 237 |

The SP19 independent dataset only contains proteins from three groups: Eukaryotes, Gram-positive, and Gram-negative bacteria. It is generated by removing proteins released in papers prior to April 2018 and proteins composed of less than 30 amino acids. Furthermore, to better ensure the independency of the SP19 dataset, CD-HIT is applied to remove redundant proteins sharing more than 20% redundancy with proteins in SiganlP5.0 database. Table 2 shows the statistics of the independent dataset.

**Table 2:** Statistics of independent dataset adopted in this study

| Dataset | Organism | SP | Total |
| --- | --- | --- | --- |
| SPDS19 | Eukaryotes | 33 | 33 |
|  | Gram-positive | 9 | 9 |
|  | Gram-negative | 3 | 3 |

### 2.2 Model Overview

Bi-directional long short-term memory (Bi-LSTM) is a special version of the recurrent neural networks, which is able to learn long-distance dependencies from input multi-channel embeddings and has been successfully used to solve signal peptide prediction problem [18]. The soft attention mechanism achieves great successes in a broad range of fields, such as image classification, text translation *etc*. Furthermore, the model architecture presented in [19] has unveiled the possibility to boost classification performance via the combination of soft attention mechanism and Bi-LSTM. To deal with these issues, we develop a novel model based on model architecture of TargetP2.0 and SignalP3L 3.0. We build up a model called USPNet to simultaneously predict signal peptide type and cleavage sites. In this case, the input sequences are the first 70 amino acids of proteins according to setting of SignalP5.0. A sequence is a multi-channel vector with each channel corresponding to a specific residue type. Figure 1 summarizes the architecture of the proposed USPNet method.

**Figure 1:**
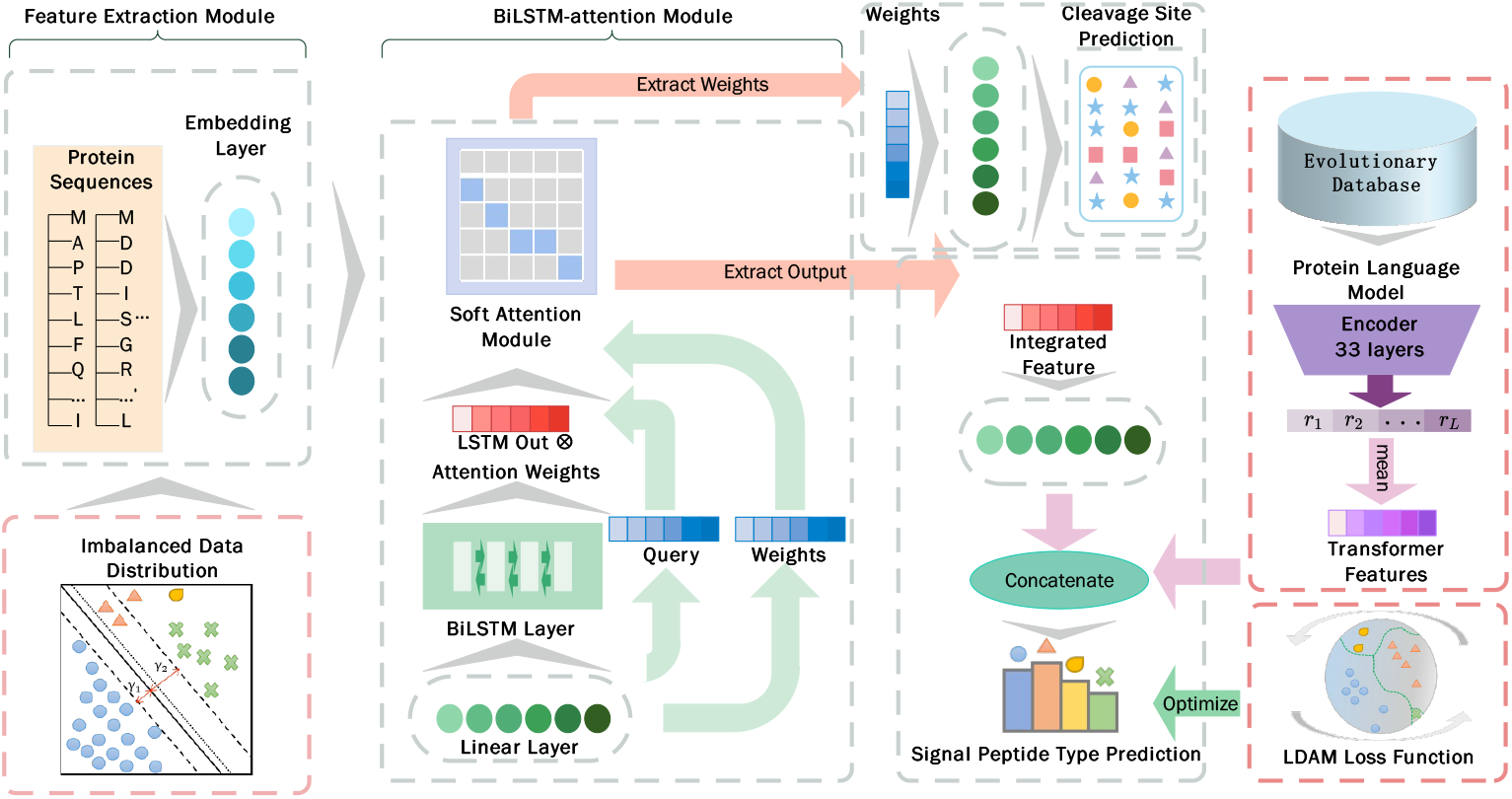
The architecture of the classifier for predicting signal peptide (SP), where L is the length of input sequence and set to be 70 in the module.

### 2.3 Model Architecture

For USPNet, the model architecture is composed of two key components: The feature extraction module and the BiLSTM-attention model to both predict signal peptide type and cleavage sites. According to SignalP5.0, we set a cut-off as 70 to sequence length of proteins, which means that each input sequence only contains 70 amino acids. Considering the number of different residue types is 20, we use an embedding layer functioning as feature extraction layer to turn input sequences into L×20-D matrix;

Then we describe the model architecture of USPNet from high level. The first part is a BiLSTM module with soft attention mechanism. The embedding vectors are firstly fed into one fully connected layer with 32 hidden units. The following procedure is to utilize a BiLSTM layer with 256 hidden units to strongly extract long-distance dependencies of sequences over forward and backward directions. In this way, we integrate information from input sequences in a higher level for later prediction by concatenating the outputs of two directions. The first four input states of the BiLSTM can be utilized to contain some group information (proved to be not necessary in later experiments). To be more specific, these states represent an one-hot encoding vector related to Eukaryotes, Grampositive, Gram-negative bacteria and Archaea. We concatenate the outputs of both directions into 512-dimensional vector for later processing:

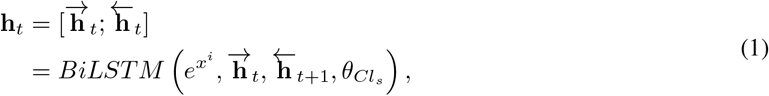

where 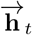 and 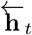 denote hidden states at time point t over forward and backward directions, respectively.

Considering that soft attention mechanism is possible to boost the performance of LSTM [20], we use a modified soft-attention-lstm module here to learn dependencies from embedding vectors:

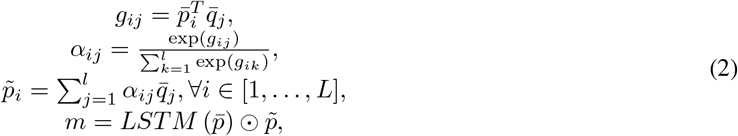

where 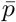 and 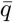 are both outputs of the first fully connected layer, which indicate key and query of the attention module respectively; *g_ij_* denotes the attention weight, which unveils the correlation between each residue of sequences; *α_ij_* is the score calculated for each hidden state in value vector according to softmax function; 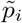 is the soft attention function computed by applying the normalized score *α_ij_* to each hidden state along the sequences; *L* is the length of input sequence, which is set to 70 by default; *m* represents the output of attention module, which is equal to the dot product of output of LSTM layer and former computed soft attention function 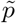.

The final output of the soft-attention-LSTM module will keep the same number of channels as its input. To better aggregate information from different feature subspaces, we develop a multi-head attention mechanism in the attention module. We conduct the above operations [Equations (2)] in each head and concatenate the multiple single-headed outputs together:

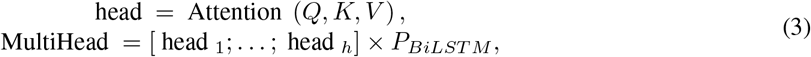

where MultiHead denotes the outputs of multi-head attention module; h represents the number of heads and *P_BiLST M_* represents output of soft-attention-LSTM layer.

Then, the outputs of this attention module will be firstly fed into two fully connected layers and used for computing a soft multi-attention matrix, which is similar to that in the model architecture of TargetP2.0. The difference is that the size of the context matrix is set to 144×9. The 9 output attention vectors correspond to 9 cleavage site types of each residue for signal peptides. The matrix is used to produce multiple representations of input sequences, with different weights over different parts of given sequences. The attention weights of the matrix are processed by two fully connected layers to directly predict the presence of cleavage sites. At the same time, the dot product of the output of attention-LSTM module and the context attention matrix is used for further processing to predict signal peptide type.

Before we summarize the outputs into logits for direct prediction, we introduce ESM embeddings here to intergrate more information. We extract 1280-dimensional embeddings from the final layer of the BERT model and use an aggregation of fully connected layers to acquire high-level information of embeddings. The processed results are concatenated with the feature matrix calculated based on context attention matrix and BiLSTM outputs. The concatenated results have 512×32+64 channels and are summarized by a fully connected layer with 256 units. Finally, the outputs are forwarded to a normalized linear projection layer and soft activation.

### 2.4 Improve Generalization

Most existing knowledge-bases related to signal peptides suffer extreme data imbalance. The number of the minority classes of signal peptide sequences is usually much smaller than that of non-signal-peptide sequences. This scenario may lead to poor generalization in low-resource data types. However, most previous models chose cross-entropy function as the objective function and ignored the data imbalance problem. There are some existing techniques solving the data imbalance problem. Cost-sensitive re-sampling and re-weighting can effectively cope with challenges brought by the data imbalance. However, for the signal peptide prediction, these methods are case-sensitive and prone to overfitting. Post-correction methods are also proposed to handle data imbalance, but the generalization improvement cannot be ensured. Inspired by the vanilla empirical risk minimization (ERM) algorithm, LDAM loss is proposed to be combined with reweighting to solve the issue.

LDAM loss focuses on correcting the cross-entropy function by introducing the margin item Δ*y* to improve generalization of classes. It suggests that a suitable margin should achieve good trade-offs between the generalization of major classes and minor classes. The class-dependent margin for multi classes is verified to have the form:

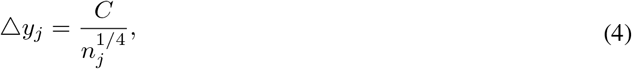

However, the margin item has changed the amplitude of logits of correct category and influenced generalization, which adds up to the difficulties of representation learning and makes model sensitive to parameter settings. Here, we introduce agent vectors for each class by applying a normalized linear layer in the final layer of classifier [21]. We normalize both the inputs and the weights of the linear projection layer to clamp the inner product of feature vectors and weight vectors into [−1, 1]. The experiments prove that the normalization can not only improve the robustness of the model, but also accelerate the convergence during the training process.

Empirically, softmax cross-entropy function with scaling factor is widely used in reinforcement learning and relevant fields. On one hand, if the scaling factor is too large, class intervals will become close to 0 and therefore influence generalization. On the other hand, a small scaling factor will lead to deviation with objective function. Here, a scaling factor is used as a hyper-parameter to ensure the final logits locate in a reasonable scope:

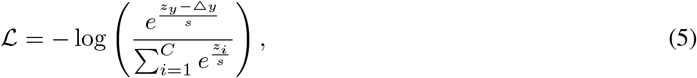

where *z_y_* denotes the logit score of groud true label; *z_i_* denotes the logit score of other labels, and s represents scaling factor. Both *z_y_* and *z_i_* are normalized as agent vectors.

### 2.5 State-of-the-art Feature Representation

Inspired by the major progress of unsupervised learning in biology, we decide to utilize the so-called ESM embeddings acquired by the ESM model to enrich the representation of USPNet. The model is trained based on 250 million protein sequences of the Uniprot database, which is large enough to support the high-capacity protein language model. And deep transformer is chosen as the architecture, because it has overperformed other recurrent architectures in many tasks. The amino acid sequences extracted from database are further divided into different fractions and given a special mask token as inputs of model in the training process. The output of neural network is the missing token of corresponding sequences.

Compared with other protein language models, the ESM model makes the best of the powerful model architecture and large evolutionary database. In addition, the unsupervised learning is good at encoding properties of protein sequences in many different scales, which contributes to the high performance of the model. It is verified that without biological signal other than sequences, the model can learn the structures of amino acids, protein sequences and evolutionary homology [16]. Furthermore, the generalization of model trained over multi-families is better than the effects based on single family. Accordingly, we believe that by adding the representation of the ESM model, the similarity of sequences within the single group can be captured and the USPNet will become more powerful to differentiate sequences from different group organisms.

## 3 Results

### 3.1 Overview

In this section, we evaluate the performance of USPNet on both classification and cleavage site prediction. The experiments can be divided into four parts: i) Overall performance comparison against existing methods; ii) Performance of organism-agnostic model; iii) The performance over SP19 independent dataset; iv) Ablation study of the effects of different losses and features.

In our experiments, all the models are trained on SignalP5.0 dataset. We applied the Adam optimizer [22] with initial learning rate of 2×10^*−*3^ and a weight decay of 1×10^*−*3^ [23]. The total number of epochs is set to be 300. All the experiments run on four V100 GPU cards with 32GB memory.

### 3.2 Evaluation criteria

For the taken metrics in the experiments, we summarize them into two components: Metrics used for evaluation of classification performance and cleavage site prediction performance. For classification, we continue to take the metrics of SignalP5.0 to test the performance of USPNet in the first part for better comparison. The two-time run pipeline is applied here. We used Matthews correlation coefficient (MCC) as measurements and only considered true and false positive and negative in sequence level:

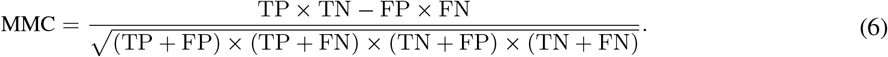

In the first run, we only take globular and/or transmembrane proteins as negative set and proteins of relevant signal peptide type as positive set (MCC1). And then, in the second run, all the rest sequences are added to the negative set (MCC2).

While for ablation study, we not only evaluate classification performance by MCC over different organism groups, but also apply overall MCC, Kappa and balanced accuracy to evaluation. This is because in ablation study, some models’ performances in minor classes are pretty close, and the overall performances will be more straightforward and intuitive to display differences. Kappa is broadly used in consistency test and measurements of multi-classification. And considering the extreme imbalance in datasets. It is not appropriate to apply accuracy to the dataset of signal peptides. Instead, we take balanced accuracy to measure the overall performance of models, which normalizes true positive and true negative predictions over the total number of samples:

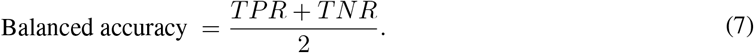

In the cleavage sites prediction part, we again follow SignalP5.0 to take precision and recall as measurements. Precision represents the fraction of correct cleavage site predictions over the total number of predicted cleavage sites, and recall represents the fraction of correct cleavage site predictions over the number of ground true labels of cleavage sites.

### 3.3 Comparison to existing methods

The USPNet model is able to predict signal peptides type and cleavage site at the same time. We compare USP-Net with some existing methods including the benchmark model, SignalP5.0, to further spotlight the advantages of USPNet model.

In this part, we collect results over different organism groups from several known baseline models. According to Table 3, Table 4, and Table 5, it is obvious that USPNet achieves a higher MMC than SignalP5.0. Over all the group categories, USPNet achieves the best performance compared with all the former proposed models.

**Table 3:** Benchmarking of Sec/SPI signal peptide detection predictions

| Method | Archaea |  | Eukaryotes | Gram-negative |  | Gram-positive |  |
| --- | --- | --- | --- | --- | --- | --- | --- |
|  | MCC1 | MCC2 | MCC | MCC1 | MCC2 | MCC1 | MCC2 |
| SignalP4.1 | n.d. | n.d. | 0.808 | 0.851 | 0.248 | 0.949 | 0.148 |
| DeepSig | n.d. | n.d. | 0.819 | 0.792 | 0.166 | 0.870 | 0.155 |
| SignalP5.0 | **0.922** | **0.917** | 0.883 | 0.860 | 0.830 | 0.922 | 0.760 |
| USPNet | **0.922** | **0.917** | **0.901** | **0.876** | **0.851** | **0.975** | **0.861** |

**Table 4:** Benchmarking of Sec/SPII signal peptide detection predictions

| Method | Archaea |  | Gram-negative |  | Gram-positive |  |
| --- | --- | --- | --- | --- | --- | --- |
|  | MCC1 | MCC2 | MCC1 | MCC2 | MCC1 | MCC2 |
| PRED-LIPO | 0.766 | 0.743 | 0.629 | 0.707 | 0.775 | 0.775 |
| LipoP | 0.836 | 0.755 | 0.791 | 0.833 | 0.798 | 0.822 |
| SignalP5.0 | **0.936** | 0.910 | **0.945** | 0.946 | 0.917 | 0.923 |
| USPNet | **0.936** | **0.940** | **0.945** | **0.948** | **0.946** | **0.938** |

**Table 5:**
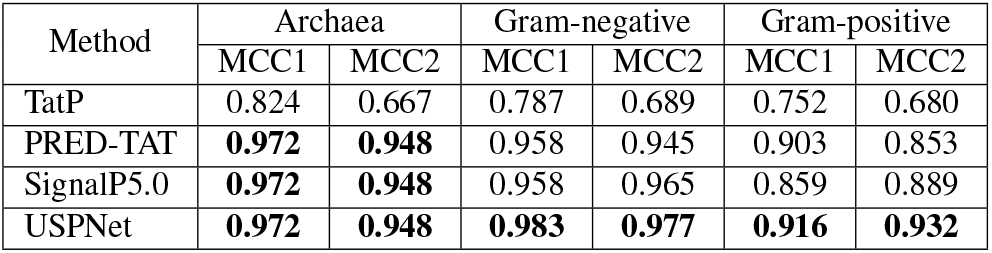
Benchmarking of Tat/SPI signal peptide detection predictions

| Method | Archaea |  | Gram-negative |  | Gram-positive |  |
| --- | --- | --- | --- | --- | --- | --- |
|  | MCC1 | MCC2 | MCC1 | MCC2 | MCC1 | MCC2 |
| TatP | 0.824 | 0.667 | 0.787 | 0.689 | 0.752 | 0.680 |
| PRED-TAT | **0.972** | **0.948** | 0.958 | 0.945 | 0.903 | 0.853 |
| SignalP5.0 | **0.972** | **0.948** | 0.958 | 0.965 | 0.859 | 0.889 |
| USPNet | **0.972** | **0.948** | **0.983** | **0.977** | **0.916** | **0.932** |

Generally, the key focus of USPNet is to produce a more generalizable classifier to classify various signal peptide types based on better generalization of different classes. But we also utilize the attention weights of context attention matrix to acquire information related to per amino acid and make decisions about cleavage sites. For fair comparison, we employ precision and recall to evaluate USPNet’s performances on cleavage site prediction against SignalP5.0.

From Figure 2, we observe that the overall performances on cleavage site predictions of USPNet are comparable to SignalP5.0. Specifically, the precision of USPNet is superior even though the recall yields lower than SignalP5.0. And unlike SignalP-3L 3.0 and other models relying on PSSM and HMM profiles to enrich the embeddings and enhance performances, USPNet is more straightforward to make predictions without evolutionary profile-based features, therefore ensuring shorter processing time.

**Figure 2:**
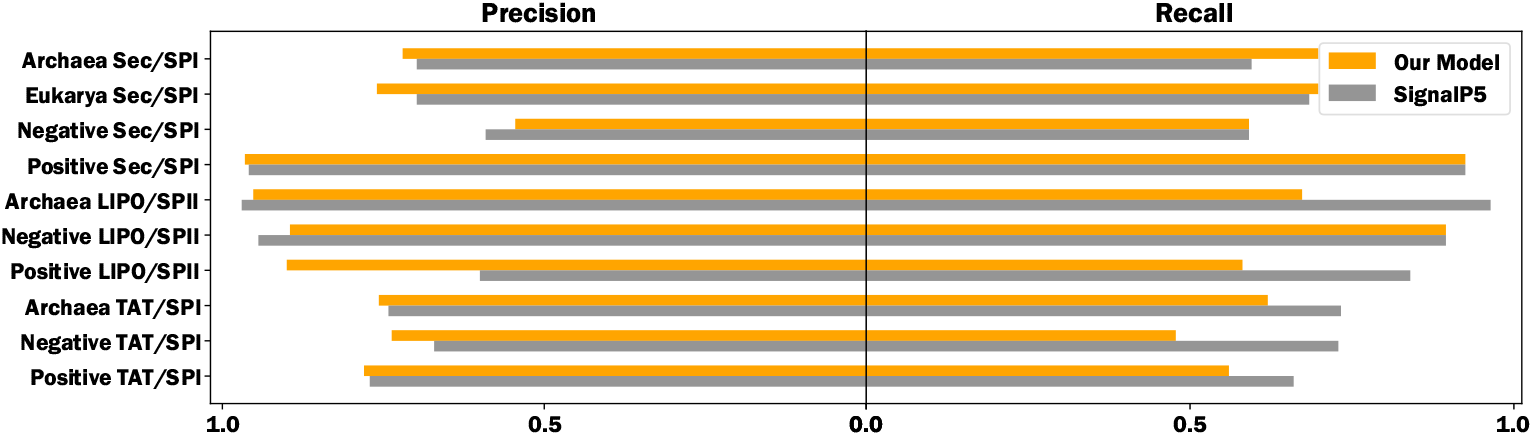
USPNet shows strong performance comparable to SignalP5.0 on cleavage site prediction.

### 3.4 Unbiased performance evaluation

The SignalP5.0 dataset contains 17270 sequences and can be considered into four parts according to four different kinds of labels. However, only 456 of them are labeled as Tat/SPI signal peptides, and 1540 are labeled as Sec/SPII signal peptides (Table 1). The sequences with major-class labels are more than ten times as the sequences with minor-class labels. Therefore, we evaluate the unbiased performance by focusing on the MCC performance over different divisions of sequences belonging to rare classes.

According to Table 4 and Table 5, it is obvious that USPNet has made an impressive promotion in the performances over data labeled as minor classes. Especially for the ground true label as Tat/SPI, performances (MCC) on sequences of gram-positive group and gram-negative increase by 6.6% and 2.6%, respectively. This accords with our assumption that the margin-based loss formulation allow the classification boundary of rare classes to be extended further and avoid overfitting in some ways. With better generalization, USPNet becomes the first unbiased multi-class SP predictor, which fit to the situations in the metagenomic research better.

### 3.5 Performance of organism-agnostic model

We have shown that USPNet is an unbiased prediction tool that outperforms other previous models in classifying signal peptide types. In this part, we will focus on comparing the organism-agnostic classification performance of USPNet against benchmark model.

Most previous proposed signal peptide prediction tools including SignalP5.0 and TargetP2.0 rely on extra group information of sequences to enrich embeddings and boost the performance. Usually, a one hot embedding vector with specific number of dimensions corresponding to group origins will be fed into the LSTM layer to produce high-level features. However, it is not realistic to produce group information for most situations in the metagenomic research. Under most conditions, we expect to accurately predict signal peptide types according to inputting amino acid sequences.

Inspired by the improvement of ability of self-supervision to learn representations based on large datasets, a high-capacity transformer protein language model is set up. The metric structure is preserved in the representation space. And by applying the protein language model to a general range of applications, the transformer is verified to have the state-of-the-art performance in differentiating raw sequences. The intuition is that the protein language model is able to project protein embeddings into a high-level feature space and represent biological variation. The representations will possibly substitute for the formerly mentioned group embedding. With the so-called Evolutionary Scale Modeling (ESM) embeddings, USPNet is turned into an organism-agnostic model to handle the universal database.

We firstly evaluate SignalP5.0’s reliability on the group information. The prediction results are based on web-servers. The benchmark dataset of SignalP5.0 can be divided into four categories: Eukarya, Archaea, Gram Positive and Gram Negative. Therefore, for each category, we will input a four-dimensional one-hot vector corresponding to the four categories. The results show that only the group information embedding in accord with ground true group type can help predictor reach promised accuracy on signal peptide prediction (Figure 3). For inputs with mismatched group information, there is an obvious decline over all the categories. The differences in MCC between the best prediction results and the worst prediction results for Eukarya, Archaea, Gram Positive and Gram Negative are 0.3243, 0.3693, 0.7066, and 0.6881, respectively. The signal peptides of minor-classes concentrate on groups of Gram Positive and Gram Negative, which suffer the most when group mismatching happens.

**Figure 3:**
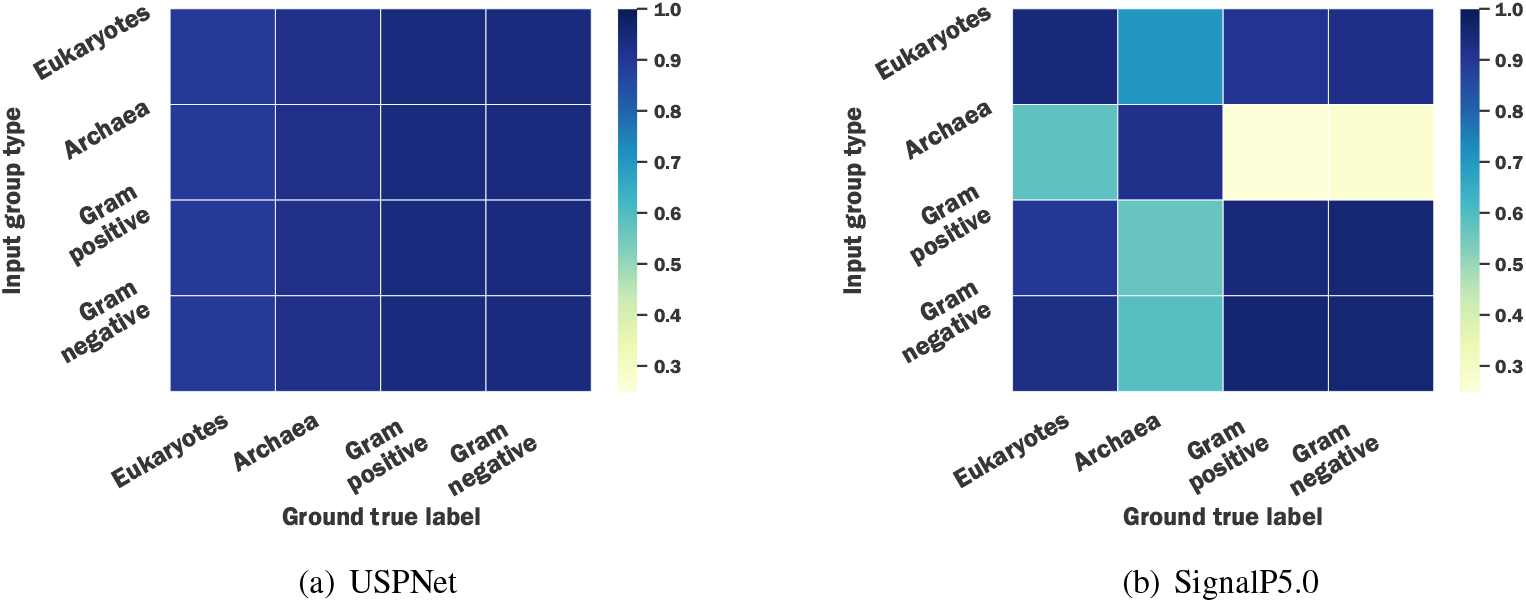
Heatmaps of performance of models with different mismatching ratios (based on MCC).

We also design test cases to input group information with different mismatching ratio, which means the proportion of the number of incorrectly matched sequences in the total number of sequences of benchmark dataset. For simplicity, unrelated group labels take an uniform distribution over the incorrect part. Under these conditions, we can clearly display to what extend SignalP5.0 is dependent on the group information. The mismatching ratios are set to 0.25, 0.375, 0.50, 0.625, and 0.75, respectively. We compare the overall performances (MCC) of USPNet and SignalP5.0 in Figure 4.

**Figure 4:**
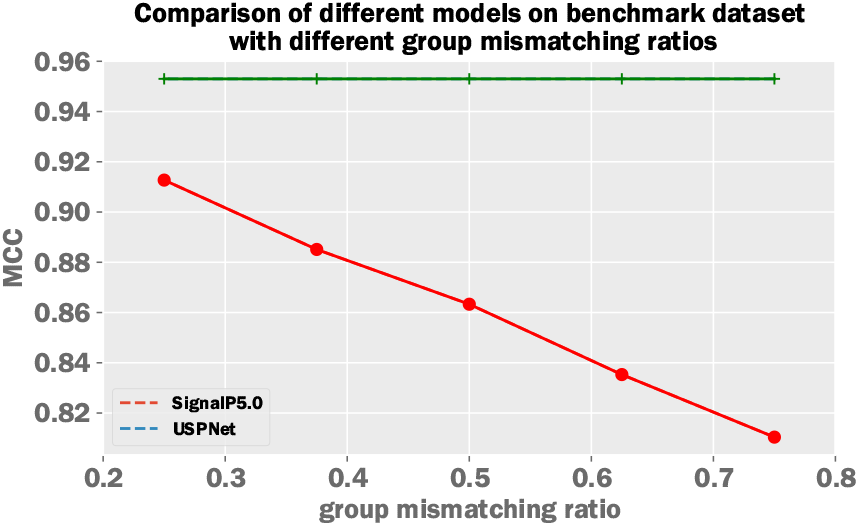
Comparison of different models with different mismatching ratios.

Even when we only set a mismatching ratio of 0.25, the SignalP5.0 model’s MCC on the benchmark dataset has decreased to 0.9127. In comparison, the performances of USPNet without group information only leads to slight decline in performance. To be more specific, when we take totally randomized group embedding vector as auxiliary inputs, the MCC performance over benchmark dataset of USPNet is 17.5% higher than SignalP5.0. This means that the USPNet is a more robust tool to make accurate predictions without the group information of organisms. From the results, we can see that USPNet is an organism-agnostic end-to-end model to directly get results based on residue-level sequences. This can be beneficial for typical cases of the metagenomic data.

### 3.6 Results on SP19 independent dataset

SignalP-3L 3.0 provides the SP19 independent dataset with strict limitation to differentiate the knowledge-base from SignalP 5.0 benchmark dataset. Accordingly, we analyse the performances of USPNet on the SP19 dataset for further verification. We collect results by feeding sequences into web-servers of several well-known signal peptide predictors: SignalP-3L 3.0 and SignalP5.0. And the results of comparison are shown in Figure 6.

There are three organism groups in total among sequences of SP19, namely 33 Eukaryotes SP sequences, 3 Gram negative SP sequences, and 9 Gram positive sequences. USPNet, SignalP-3L 3.0, and SignalP5.0 reach equal performances on Eukaryotes SP sequences. All of them can accurately predict the appearance of signal peptides given amino acid sequences. However, for Gram negative and Gram positive, only USPNet and SignalP-3L 3.0 can acquire the best performances. It further unveils the advantages of USPNet in learning features of data with minor-class labels.

### 3.7 Ablation study

In this part, we will discuss performances of models with different loss functions and embeddings to deeply look into the reasons why USPNet become more universal and organism-agnostic. We will compare LDAM loss with other well known loss functions. Also, we will evaluate the differences in performances between models with and without ESM embeddings.

We firstly trained the USPNet model with three kinds of loss functions: cross-entropy function, focal loss and LDAM loss [23]. Cross-entropy loss is widely used for the training of deep-learning-based signal peptide predictors in the last few years. And focal loss has achieved great successes in many fields such as image classification and entity recognition. We take these two loss functions here for better comparability to display the advantages of LDAM loss. The hyperparameters of loss functions are chosen manually according to multiple runs.

By analyzing the results of experiments (Figure 5), it is clear that LDAM loss allows for the maximum gains in improving the accuracy of predictions. With the same model architecture and ESM embedding, the LDAM ESM model trained based on LDAM loss reaches the MCC of 0.957, which is 1.8% higher than the Focal loss and 2.6% higher than cross-entropy loss, respectively. In the experiments, we also try to apply the deferred re-balancing training procedure proposed with the LDAM Loss. Reweighting is considered to be an orthogonal modification of the loss function. According to this point, Class-Balanced Loss, which is based on adding reweighting according to the capacity of effective sample space, is therefore introduced after running for a specific number of epochs during training to boost the performance [24]. However, in our experiments, when training the signal peptide predictor, the so-called LDAM-DRW algorithm cannot help gain benefits. Instead, adding class-balanced reweighting from the start will help the model make further progress. Therefore, we propose to directly apply the combination of LDAM Loss and class-balanced loss, called CB-LDAM loss, during the training process of USPNet.

**Figure 5:**
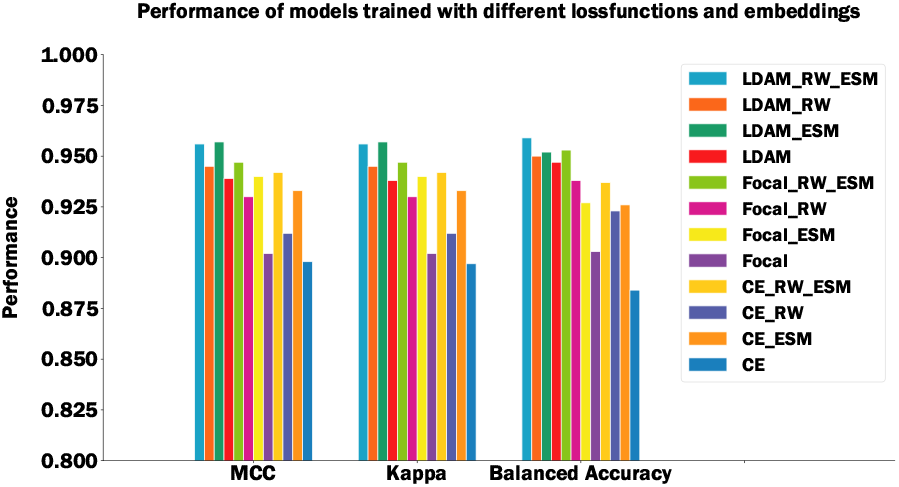
Performance of models trained with different loss functions and embeddings.

**Figure 6:**
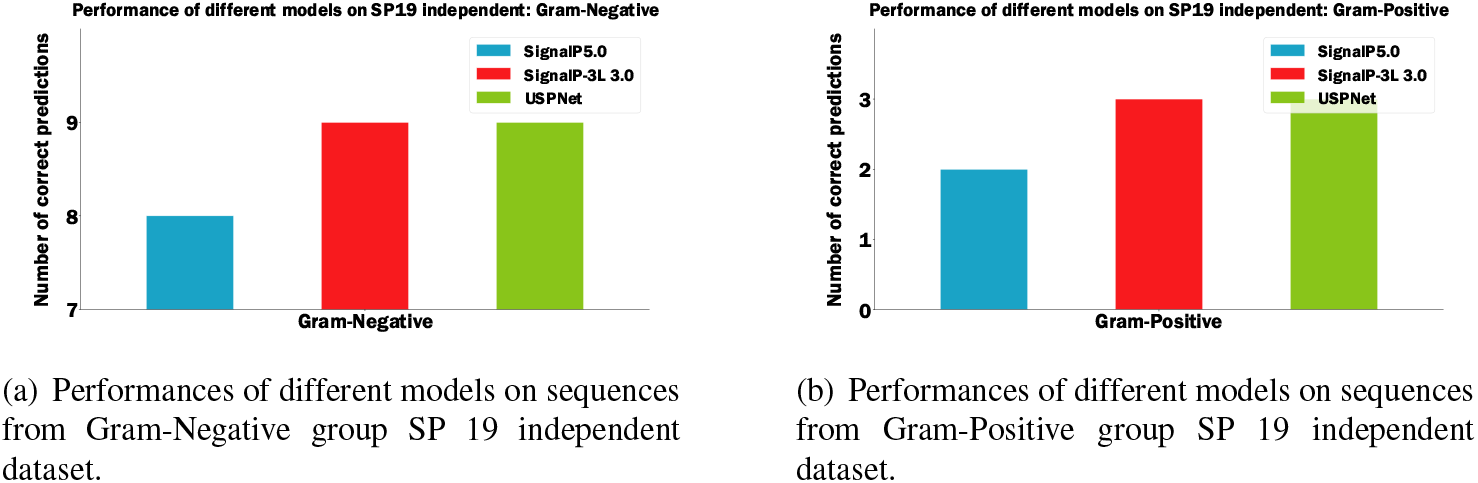
Performance of different models on SP19 independent dataset.

Except for baseline loss functions, we used focal loss and cross entropy loss with reweighting for comparison. We manually give different weights according to the frequency of appearance of different signal peptide labels to enhance generalization. And these manual reweighting factors are acquired based on the results of multiple runs. For models using ESM embeddings, we use one, two, and three, linear layers respectively to integrate information from original ESM embedding. The results of experiments suggest that we get the best results when we use two layers. And the reason for performance of decline may be overfitting. It is clear that by adding the ESM embeddings, the model can acquire continuous improvement by intergrating information from state-of-the-art embeddings.

## 4 Conclusion and Future work

In this work, we focus on the existing long-tail problem in signal peptide prediction and apply LDAM loss to solve the problem, which is orthogonal to most deep-learning-based signal peptide predictors. Based on LDAM loss, we propose to combine class-balanced loss and LDAM loss together as so-called CB-LDAM loss to enhance the generalization of boundaries of minor-classes and major-classes. Furthermore, considering most existing models’ dependence on group information, we propose to introduce ESM embeddings to achieve an end-to-end organism-agnostic model. Finally, we present a new model architecture called USPNet. The attention-based BiLSTM model allows for maximizing the possibilities to integrate and extract relationships among different positions of inputting sequences and helps boost performance. The experiment results have verified that the integration of all these techniques can achieve better performance in signal peptide prediction compared with state-of-the-art methods. Our method can be potentially useful for other similar protein function prediction problems [25, 26, 27, 28] and data imbalance tasks in computational biology [25, 29, 30].

In the future, we will apply LDAM loss and ESM embeddings can be transferred to more general fields of bioinformatics, because the data imbalance and object—dependence is very common in biological prediction problems. In addition, we will explore the performance of the proposed USPNet on more datasets and other relevant problems to evaluate the robustness of USPNet. Finally, meta-learning and multi-task learning [31], which have been successfully applied for many data-imbalance and hyperparameter-sensitive problems, will be introduced to continuously improve the robustness of the model.

## Notes

### Competing Interest Statement

The authors have declared no competing interest.

### Summary of Updates

Typo and grammar erros fixed.

## References

[1] Heijne, G. V. The signal peptide. The Journal of Membrane Biology 115, 195–201 (1990).

[2] Savojardo, C., Martelli, P. L., Fariselli, P. & Casadio, R. Deepsig: deep learning improves signal peptide detection in proteins. Bioinformatics 10 (2017).

[3] Heijne, G. V. Patterns of amino acids near signal-sequence cleavage sites. European Journal of Biochemistry 133(1983).

[4] Nielsen, H. & Krogh, A. Prediction of signal peptides and signal anchors by a hidden markov model. Intelligent Systems for Molecular Biology 6, 122–130 (1998).

[5] Henrik, N., Søren, B. & Gunnar, V. H. Machine learning approaches for the prediction of signal peptides and other protein sorting signals. Protein Engineering 3 (1999).

[6] L., K., Krogh, A. & Sonnhammer, E. A combined transmembrane topology and signal peptide prediction method. Journal of Molecular Biology 338, 1027–1036 (2004).

[7] Bendtsen, J. D., Nielsen, H., Von Heijne, G. & Brunak, S. Improved prediction of signal peptides: Signalp 3.0. Journal of molecular biology 340, 783–795 (2004).

[8] Reynolds, S. M., Käll, L., Riffle, M. E., Bilmes, J. A. & Noble, W. S. Transmembrane topology and signal peptide prediction using dynamic bayesian networks. PLoS computational biology 4, e1000213 (2008).

[9] Viklund, H., Bernsel, A., Skwark, M. & Elofsson, A. Spoctopus: a combined predictor of signal peptides and membrane protein topology. Bioinformatics 24, 2928–2929 (2008).

[10] Bagos, P. G., Nikolaou, E. P., Liakopoulos, T. D. & Tsirigos, K. D. Combined prediction of tat and sec signal peptides with hidden markov models. Bioinformatics 26, 2811–2817 (2010).

[11] Frank, K. & Sippl, M. J. High-performance signal peptide prediction based on sequence alignment techniques. Bioinformatics 24, 2172–2176 (2008).

[12] Petersen, T. et al. Petersen tn, brunak s, von heijne g, nielsen h signalp 4.0: discriminating signal peptides from transmembrane regions. (2011).

[13] Savojardo, C., Martelli, P. L., Fariselli, P. & Casadio, R. Deepsig: deep learning improves signal peptide detection in proteins. Bioinformatics 10 (2017).

[14] Armenteros, J. J. A. et al. Signalp 5.0 improves signal peptide predictions using deep neural networks. Nature biotechnology 37, 420–423 (2019).

[15] Cao, K., Wei, C., Gaidon, A., Arechiga, N. & Ma, T. Learning imbalanced datasets with label-distribution-aware margin loss. arXiv preprint arXiv:1906.07413 (2019).

[16] Rives, A. et al. Biological structure and function emerge from scaling unsupervised learning to 250 million protein sequences. Proceedings of the National Academy of Sciences 118(2021).

[17] Consortium, U. et al. Uniprot: the universal protein knowledgebase. Nucleic acids research 46, 2699 (2018).

[18] Thireou, T. & Reczko, M. Bidirectional long short-term memory networks for predicting the subcellular localization of eukaryotic proteins. IEEE/ACM transactions on computational biology and bioinformatics 4, 441–446 (2007).

[19] Zhang, W.-X., Pan, X. & Shen, H.-B. Signal-3l 3.0: improving signal peptide prediction through combining attention deep learning with window-based scoring. Journal of Chemical Information and Modeling 60, 3679–3686 (2020).

[20] Chen, Q. et al. Enhanced lstm for natural language inference. arXiv preprint arXiv:1609.06038 (2016).

[21] Wang, F., Xiang, X., Cheng, J. & Yuille, A. L. Normface: L2 hypersphere embedding for face verification. In Proceedings of the 25th ACM international conference on Multimedia, 1041–1049 (2017).

[22] Kingma, D. P. & Ba, J. Adam: A method for stochastic optimization. arXiv preprint arXiv:1412.6980 (2014).

[23] Lin, T.-Y., Goyal, P., Girshick, R., He, K. & Dollár, P. Focal loss for dense object detection. In Proceedings of the IEEE international conference on computer vision, 2980–2988 (2017).

[24] Cui, Y., Jia, M., Lin, T.-Y., Song, Y. & Belongie, S. Class-balanced loss based on effective number of samples. In Proceedings of the IEEE/CVF conference on computer vision and pattern recognition, 9268–9277 (2019).

[25] Li, Y. et al. Deepre: sequence-based enzyme ec number prediction by deep learning. Bioinformatics 34, 760–769 (2018).

[26] Zou, Z., Tian, S., Gao, X. & Li, Y. mldeepre: Multi-functional enzyme function prediction with hierarchical multi-label deep learning. Frontiers in Genetics 9, 714 (2019).

[27] Lam, J. H. et al. A deep learning framework to predict binding preference of rna constituents on protein surface. Nature communications 10, 1–13 (2019).

[28] Wei, J., Chen, S., Zong, L., Gao, X. & Li, Y. Protein-rna interaction prediction with deep learning: Structure matters. arXiv preprint arXiv:2107.12243 (2021).

[29] Umarov, R., Kuwahara, H., Li, Y., Gao, X. & Solovyev, V. Promoter analysis and prediction in the human genome using sequence-based deep learning models. Bioinformatics (2019).

[30] Chen, X., Li, Y., Umarov, R., Gao, X. & Song, L. Rna secondary structure prediction by learning unrolled algorithms. In International Conference on Learning Representations 2020 (2020).

[31] Li, Y. et al. Hmd-arg: hierarchical multi-task deep learning for annotating antibiotic resistance genes. Microbiome 9, 1–12 (2021).

